# Microglial morphology reflects cognitive status in the aging rat brain

**DOI:** 10.1101/2025.10.14.682389

**Authors:** Sarah J. Myers, Austyn D. Roseborough, Cristian X. Bayona, Carlangelo Carrese, Brian L. Allman, Shawn N. Whitehead

**Author notes:** Correspondence: Shawn Whitehead, PhD Department of Anatomy and Cell Biology 458 Medical Sciences Building Western University London, Ontario N6A 3K7.

## Abstract

Age-related cognitive decline affects millions of individuals worldwide, but the cellular mechanisms underlying this decline remain incompletely understood. Microglia undergo significant changes with aging, including alterations in morphology, that may reflect or contribute to cognitive dysfunction. However, the relationship between specific microglial morphologies and cognitive performance in relevant brain regions remains poorly understood. To address this, we evaluated the relationship between morphology-based microglial phenotypes and cognitive performance across domains affected by aging. Microglial morphology was analyzed in four cognitive brain regions of male and female 3-, 9-, and 15-month-old rats and features were subjected to hierarchical clustering on principal components to identify microglial subtypes. Rats underwent cognitive testing using a radial arm water maze and a T-maze set-shifting task to assess spatial working and reference memory, striatal-based learning, and cognitive flexibility. We observed age-related cognitive impairments alongside region-specific changes in microglial morphotype abundance. Importantly, the relative abundance of distinct microglial clusters correlated with cognitive performance in functionally relevant brain regions including the prefrontal cortex, the orbitofrontal cortex, and the hippocampus. Taken together, these findings highlight the utility of morphological profiling in capturing microglial heterogeneity and suggest that morphological changes may reflect or contribute to cognitive decline during aging.

## 1. Introduction

The worldwide population of people aged 65 or older is expected to more than double to over 1.5 billion by 2050^1^. Given that age is the number one risk factor for dementia, understanding how aging affects the brain is critically important for identifying mechanisms that contribute to cognitive vulnerability in our aging population^2^. While dementia is not a part of normal aging, varying degrees of age-related decline in domains such as learning, memory, and executive function have been well-documented in both clinical and experimental studies^3–6^. These cognitive domains rely on brain regions such as the prefrontal cortex, the orbitofrontal cortex, the hippocampus, and the striatum, all of which are vulnerable to age-related structural and functional changes clinically^7–9^. While the mechanisms underlying cognitive aging are not yet well understood, microglia have emerged as essential mediators of brain homeostasis and immune response. Under normal conditions, microglia support neural function through debris clearance, myelin maintenance, and synaptic surveillance^10–12^. However, microglia are highly plastic cells that can rapidly shift their phenotype in response to pathological stimuli, injury, or aging, resulting in altered surveillance, phagocytic, and synaptic regulatory functions^13,14^. Nevertheless, how microglial phenotypes change across different brain regions during normal aging, and whether these changes correlate with cognitive decline, has not been comprehensively investigated.

Although microglial phenotyping has primarily been conducted in disease states, clinical and experimental evidence suggests microglia undergo age-related changes in gene expression, cell surface markers, functions, and morphology. Histological studies have shown that aging is associated with increased prevalence of reactive microglia that express markers such as MHC-II, CD68, and CD11c^4,15–18^. Transcriptomic and proteomic analyses further support these findings, identifying distinct age-associated microglial phenotypes that are also modulated by biological sex^10,19–22^. These molecular changes are accompanied by alterations in microglial function, including impaired debris clearance and increased pro-inflammatory signaling^23–25^. Aged microglia have also been described with unique morphologies, including dystrophic microglia, which are characterized by reduced ramification complexity and fragmented or beaded processes^26,27^. The sustained loss of microglial processes, in contrast to their normal dynamic remodeling, may indicate a dysfunctional phenotype with reduced ability to sense and respond to environmental cues^11,28^. Given that aging disproportionately affects certain cognitive brain regions, characterizing age-related changes in microglial morphology in these areas may provide important insight into the relationship between microglial phenotype and cognitive decline.

Due to their critical roles in helping maintain the neural environment, microglia are increasingly recognized as potential contributors to cognitive decline across aging and neurodegeneration. In both human postmortem tissue and transgenic Alzheimer’s disease (AD) rodent models, elevated levels of reactive microglial markers, such as CD68, MHC-II and TREM2, have been linked to poorer cognitive performance^29–31^. Evidence from animal models also suggests that microglial reactivity in normal aging may similarly associate with impairment across multiple cognitive domains^20,32,33^. However, much of this work has relied on immunohistochemical markers and transcriptomic profiling, with structural changes remaining underexplored. Microglial morphology may reflect functional status^34^, and therefore offer insight into how microglia during aging contribute to cognitive decline in vulnerable brain regions.

Despite growing interest in microglial phenotyping, most preclinical research has focused on transgenic neurodegeneration models, leaving normal aging largely unexplored. Here, we addressed this gap using a comprehensive approach combining microglia morphological profiling alongside multidomain cognitive testing in normally aging rats. We assessed cognitive performance across spatial reference memory, spatial working memory, striatal-based learning, and cognitive flexibility using radial arm water maze and T-maze set-shifting tasks. To capture microglial heterogeneity, we applied hierarchical clustering on principal components to morphological features of Iba1^+^ cells across four cognitive brain regions (prefrontal cortex, orbitofrontal cortex, hippocampus, and dorsal striatum) in 3-, 9-, and 15-month-old male and female rats. This unsupervised approach revealed distinct microglial morphotypes whose regional abundance associated with performance in domain-relevant cognitive tasks: prefrontal cortex microglia with executive function, orbitofrontal cortex microglia with cognitive flexibility, and hippocampal microglia with spatial reference memory.

## 2. Materials and methods

### 2.1 Animals

All rats included in this study were housed in facilities maintained by Western University Animal Care and Veterinary Services on a 12-hour light/dark cycle with *ad libitum* access to food and water. Male and female wildtype Fischer 344 rats were bred and aged in-house to 3-, 9-, or 15-months-old (*n* = 10 males and 10 females per age group). The animal ethics and procedures used in this study were approved by the Animal Care Committee at Western University (protocol number 2022-1367).

### 2.2 Radial arm water maze

#### Setup

We employed a 4/8 radial arm water maze (RAWM) task adapted from Hyde et al.^35^ to assess spatial working and reference memory. In a dimly lit room, a circular tank (144 cm) was filled with room temperature water (22-23 °C) and dyed black with non-toxic acrylic paint. An 8-arm steel maze inset (height: 50.8 cm; arm length: 63.5 cm measured from centre of maze; arm width: 15.24 cm) was placed inside of the tank (Maze Engineers). The protocol occurred over 17 consecutive days, where rats underwent 1 day of habituation, 1 day of training, and 15 days of testing. Rats were tested at the same time each day and brought into the room to acclimate ∼30 mins before beginning.

#### Habituation

On the first day, the rats were habituated to the maze over six trials (∼30 s intertrial intervals) which trained them to navigate from the start arm to hidden escape platforms (3 cm below water surface) located at the ends of two of the arms. The tank was enclosed by curtains without the presence of spatial cues. For all six trials, rats swam to locate a platform from various locations and remained on the platform for 30 s before removal from the tank. For trials 1 and 2, the rats were placed in an enclosed maze arm, opposite the hidden platform. Trials 3 and 4 progressed to the rats being placed in the centre of the maze with all but one arm closed off. Finally, rats were placed facing the back of the start arm for trials 5 and 6 and allowed to swim to the open arm with a hidden platform. If rats failed to find the platform within 60 s, they were cued to the platform location.

#### Training

The next day, four visual cues were placed in a square around the maze (yellow cross, white star, orange rectangle, and green triangle) and all maze arms were opened. A given animal had four platforms to find over four training trials (∼30 s intertrial intervals). A trial consisted of the animal being placed facing the back of the start arm, swimming to find a platform and remaining on the platform for 30 s to observe the surrounding visual cues. The arm of the located platform was closed off for the next trial, and this was repeated until all four platforms were located. If rats failed to find a platform within 120 s they were guided to the nearest available platform.

#### Testing

Following training, rats were given four testing trials per day for fifteen days with the same platform locations. Testing followed the same protocol as training except located platforms were removed from the tank rather than blocked off. For rats to perform well at this task they needed to 1) avoid entering arms that never contained platforms (reference memory errors), 2) use a win-shift strategy to avoid re-entering arms that previously contained platforms within that session (working memory correct errors), and 3) avoid repeat entries into arms that never contained a platform (working memory incorrect errors). All trials were recorded using ANYmaze tracking software with a webcam (C930e; Logitech) mounted on the ceiling above the water tank. Arm entries were registered as the midpoint of a rat crossing into a given arm and were scored as errors based on the requirements above, working memory correct and incorrect errors were collapsed together for a single metric.

### 2.3 T-Maze Set-Shifting

#### Setup

We employed a water-based T-maze set-shifting task in which rats were tested in a series of visual and directional discrimination tasks over several days to evaluate striatal-based learning and cognitive flexibility. The same maze setup was used as described in section 2.2 with all but three arms blocked off to form the T-maze.

#### Visual cue discrimination (VCD)

Rats were given 12 trials per day to associate the illuminated light with the escape platform. The light/platform location pseudo-randomly alternated between sides across the 12 trials. Rats were placed in the tub facing the back of the start arm and allowed to swim to find the platform. Rats were removed from the platform and allowed to rest for ∼30 s before beginning the next trial. This portion of the task was complete when a rat was correct on at least 10 out of 12 responses.

#### Response discrimination (RD)

One hour after reaching criterion in VCD, rats were given 12 trials in which the platform remained on one side, but the light continued to pseudo-randomly alternate. This was repeated each day until a rat was correct on at least 10 out of 12 responses.

#### Reversal learning (RL)

One hour after reaching criterion in RD, rats were given 12 trials in which the platform location switched to the side opposite that of RD, but the light continued to pseudo-randomly alternate. This was repeated each day until a rat was correct on at least 10 out of 12 responses.

#### Metrics

All trials were recorded using ANYmaze tracking software with a webcam (C930e; Logitech) mounted on the ceiling above the water tank. An error was registered as the midpoint of a rat crossing into an incorrect arm. Performance in each phase was assessed by total first choice errors committed before completion of the task and number of trials taken to reach a 10 out of 12 rolling average. In the case of RD, number of trials refers to incongruent trials in which the light was on the side opposite the platform.

### 2.4 Euthanasia and brain collection

Rats were euthanized at 3-, 9-, or 15-months of age by intraperitoneal injection of pentobarbital (Euthanyl, Bimeda Animal Health Inc) and transcardial perfusion using 180 mL of 0.01 M PBS and 300 mL of 4% paraformaldehyde (PFA). Brains were collected and stored in 4% PFA at 4°C, 24 hours later, brains were moved to 30% sucrose and stored at 4 °C for at least 36 hours prior to sectioning.

### 2.5 Immunohistochemistry

Brains were cut using a cryostat (CryoStar NX50, Thermo Fisher Scientific) into 30 µm sections and stored in cryoprotectant at -20 °C prior to staining. A standard diaminobenzidine (DAB) staining protocol was followed on free-floating sections at Bregma levels +3.00, +2.00, and - 3.00. Tissue was first washed in 0.01 M PBS for 6 x 10 mins. Endogenous peroxidases were blocked via incubation in 1% H_2_O_2_ (Thermo Fisher Scientific) for 10 mins and H_2_O_2_ was washed off in 3 x 5 min 0.01 M PBS washes. Sections were then blocked with 0.2% Triton-X-100 (Sigma Aldrich) in 0.01 M PBS and 2% goat serum for 1 hour at room temperature. Overnight, tissue was incubated with primary goat anti-Iba1 (1:1000; Wako Chemicals 019-19741) in blocking solution at 4 °C. The following day, sections were rinsed in 0.01 M PBS for 3 x 5 mins prior to incubation with goat anti-rabbit biotinylated secondary antibody (1:500; Invitrogen) in blocking solution for 1 hour at room temperature. Tissue was processed using an Avidin-Biotin Complex kit (Thermo Fisher Scientific) for 1 hour at room temperature and developed using 0.05% DAB (Sigma-Aldrich), with 3 x 5 min 0.01 M PBS washes between each step. Stained tissue sections were mounted onto slides using 0.3% gelatin and airdried overnight. Lastly, the slides were dehydrated using progressive concentrations of ethanol and xylene prior to cover slipping using Depex mounting medium (Electron Microscopy Sciences).

### 2.6 Microscopy

Images were taken using an upright brightfield microscope (Nikon Eclipse Ni-E, Nikon DS Fi3 colour camera, NIS Elements Imaging) by experimenters blinded to the experimental groups. Using the 20 x objective lens and 1 µm z-step distance, z-stacks were captured for a total of 4 images per region of interest (ROI) (2 in the left hemisphere and 2 in the right hemisphere) per animal. ROIs included the prelimbic region of the prefrontal cortex (PFC, Bregma +3.00 mm), the lateral region of the orbitofrontal cortex (OFC, Bregma +3.00), the CA1 region of the hippocampus (HPC, Bregma -3.00 mm), and the dorsal striatum (STR, Bregma +2.00). White balance was automated using an off-tissue reference point and settings for light intensity, exposure, and aperture were kept consistent across images.

### 2.7 Image analysis

#### Microglial morphology

All image analysis was performed by experimenters blinded to the experimental groups. Images were preprocessed using ImageJ, briefly, Z-stack images were converted to 8-bit, inverted, and the background was subtracted using a 50-pixel rolling ball radius. Images were further processed to remove noise, and image brightness was adjusted for optimal visualization and kept consistent across all images. Next, we used 3DMorph, a semi-automatic MATLAB-based program, to reconstruct three-dimensional microglia and quantify morphological metrics^36^. Using 3DMorph, cells were defined with manually outlined parameters that exclude small objects (fragmented branches) and segment large objects (multiple merged cells). The 3DMorph script then automatically skeletonized the cells and produced an output of morphology metrics, including cell territorial volume (volume of a 3D polygon around the cell’s external points), cell volume (number of voxels multiplied by the scale), ramification index (ratio of the cell’s territory to the cell’s volume), number of endpoints, number of branchpoints, average branch length, maximum branch length, and minimum branch length. Each skeletonized cell was compared with its original image to confirm accurate segmentation, cells that were split incorrectly were removed from all analyses.

#### Cell density

Cell density was assessed using the Fiji ImageJ software^37^ cell counter plugin in which Iba1^+^ cells were manually selected in each 20 x image. The number of Iba1^+^ cells was averaged across the 4 images in each region and divided by the area to obtain an Iba1^+^ cell density in each region of interest per animal.

### 2.8 Principal components analysis and hierarchical clustering

Eight microglial morphology variables (cell territory, cell volume, ramification index, endpoints, branchpoints, average branch length, maximum branch length, and minimum branch length) from all cells were Z-transformed to scale and center the data. To reduce dimensionality, principal components analysis (PCA) was applied. Variables were assessed for inclusion via a correlation matrix which confirmed that each variable had an r value ≥ 3 with at least one other variable. PCA on 8 morphological variables revealed two PCS with an eigenvalue >1 which were retained for clustering. Agglomerative hierarchical clustering on PC1 and PC2 was then performed using Ward’s method^38^. The number of clusters was determined using the Thorndike procedure^39^, in which the average within-cluster distances were plotted for different cluster numbers and the flattening of the curve represented the appropriate number of clusters. An animals’ proportion of cells in each cluster was square-root transformed and then the Z-score was calculated to show relative abundance of each cluster in each brain region.

### 2.9 Statistical analyses

Statistical analyses were conducted using SPSS (IBM Corporation) and GraphPad Prism. Depending on the comparison, one- or two-way analysis of variances (ANOVA) were performed with a significance value of *p* = 0.05 and when required, Bonferroni’s *post-hoc* correction was used. When Mauchly’s test of sphericity was violated in the mixed ANOVAs, the Greenhouse-Geisser correction was applied. Normality was assessed using the Shapiro-Wilk test and when violated, the Kruskal-Wallis one-way ANOVA with Dunn’s *post-hoc* correction was used. All data are presented as mean values with error bars indicating standard error of the mean (SEM). Methodology schematics were generated using BioRender (Biorender.com) and graphs were created using GraphPad Prism.

## 3. Results

### 3.1 Iba1^+^ microglia cell numbers were largely stable across age, except in the female striatum

To establish whether morphological changes occurred independently of cell number alterations, we first quantified microglial density across age groups and brain regions. Microglia were visualized using Iba1 immunohistochemistry in 3-, 9-, and 15-month-old male and female rats. Cells were counted in 4 regions of interest that were chosen based on their importance for cognitive performance, including the prelimbic region of the prefrontal cortex (Fig. 1A; PFC), the lateral region of the orbitofrontal cortex (Fig. 1D; OFC), the CA1 region of the hippocampus (Fig. 1G; HPC), and the dorsal striatum (Fig. 1J; STR). There were no significant age-related differences in the prefrontal cortex, (Fig. 1C), orbitofrontal cortex (Fig. 1F), or hippocampus (Fig. 1I), but there was a decrease in microglial cell density, indicating decreased numbers of microglia, in the striatum of 15-month-old females compared to 3- and 9-month-old females (Fig. 1L; One-way ANOVA: F_(2, 12)_ = 8.198, *p* = 0.006, η^2^ = 0.577; 3 vs 15: *p*_bonf_ = 0.008; 9 vs 15: *p*_bonf_ = 0.022).

**Figure 1.**
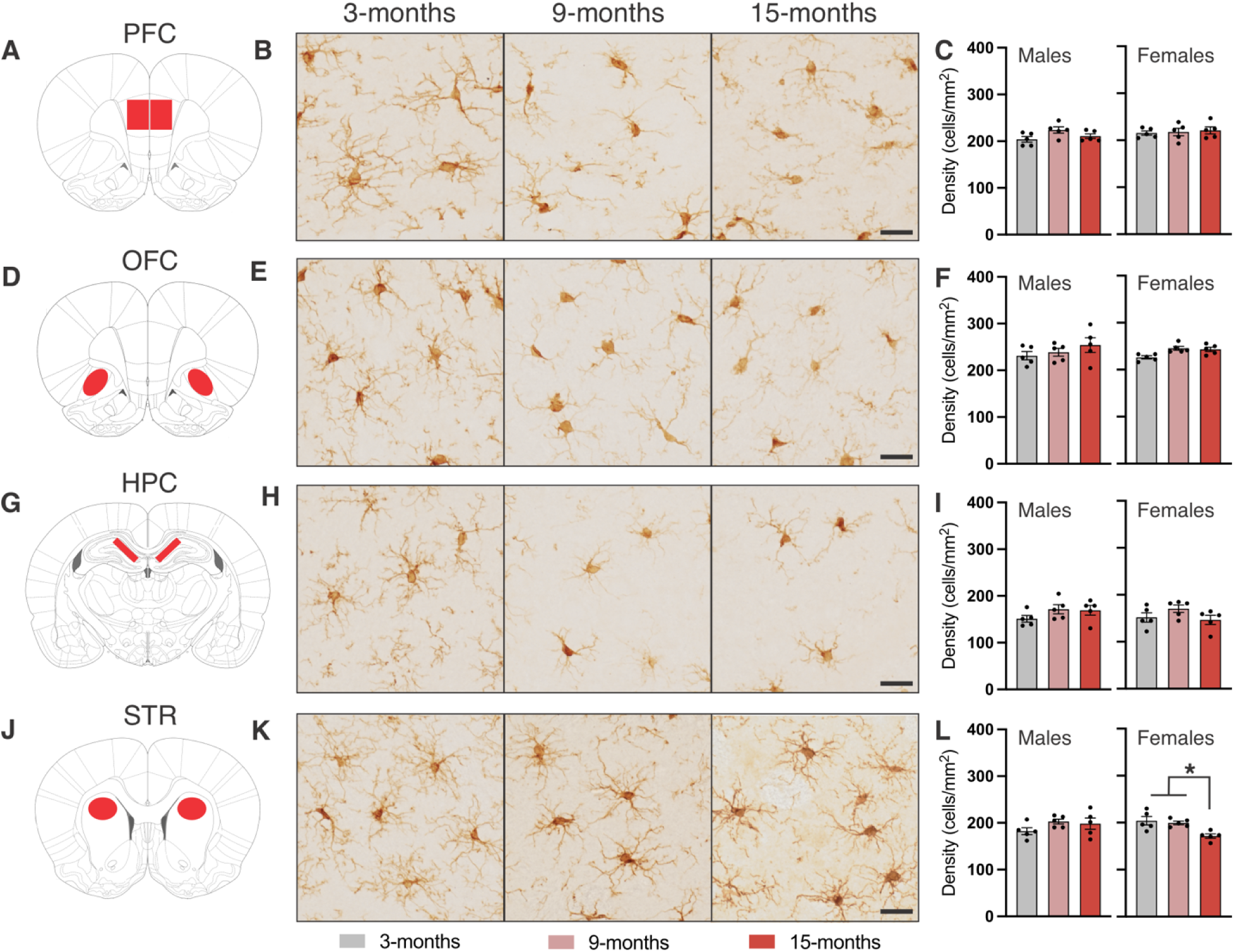
Unaltered microglial cell densities across aging, except for the female striatum. A, D, G, J) Coronal section and outline for the prefrontal cortex (PFC), orbitofrontal cortex (OFC), hippocampus (HPC), and striatum (STR). B, E, H, K) Representative 20 x images of Iba1^+^ microglia in 3-, 9-, and 15-month WT rats in each region of interest. C, F, I, L) Quantification of Iba1 cell density in male and female 3-, 9-, and 15-month WT rats in each region of interest. Scale bar indicates 25 µm. * Indicates statistical significance (*p* < 0.05) measured using a one-way ANOVA. Data represent the group mean ± SEM. *n* = 5 males and 5 females per group.

### 3.2 Age-related reduction in microglial ramification index

To quantify age-related changes in microglial morphology across our regions of interest, we used 3DMorph, a semi-automatic MATLAB-based program that generates multiple morphometric parameters from z-stack images through automated thresholding, segmentation, and skeletonization (Fig. 2A)^36^. We initially focused on ramification index (RI), a ratio of cell territory to cell volume, with higher values indicating more ramified morphology and lower values reflecting more ameboid forms. Age-related changes in RI were assessed in male and female rats across all four brain regions. We show that there was a significant reduction in RI in both male and female 9- and 15-month-old rats compared to 3-month-old rats in the prefrontal cortex (Fig. 2B; Kruskal-Wallis One-Way: Male: H_(2)_ = 152.1, *p* < .0001, η^2^ = 0.21; 3 vs 9: *p*_dunnv_ < .0001; 3 vs 15: *p_dunn_* < .0001; Female: H_(2)_ = 88.52, *p* < .0001, η^2^ = 0.11; 3 vs 9: *p*_dunn_ < .0001; 3 vs 15: *p_dunn_* < .0001) and hippocampus (Fig. 2D; Kruskal-Wallis One-Way: Male: H_(2)_ = 79.89, *p* < .0001, η^2^ = 0.14; 3 vs 9: *p*_dunn_ < .0001; 3 vs 15: *p_dunn_* < .0001; Female: H_(2)_ = 74.81, *p* < .0001, η^2^ = 0.13; 3 vs 9: *p*_dunn_ < .0001; 3 vs 15: *p_dunn_* < .0001). Similarly, RI was reduced in the orbitofrontal cortex of 3-, and 9-month-old male rats, but was only reduced at 9-months in female rats (Fig. 2C; Kruskal-Wallis One-Way: Male: H_(2)_ = 128.1, *p* < .0001, η^2^ = 0.15; 3 vs 9: *p*_dunn_ < .0001; 3 vs 15: *p_dunn_* < .0001; Female: H_(2)_ = 10.28, *p* < 0.0058, η^2^ = 0.0099; 3 vs 9: *p*_dunn_ < 0.004). Lastly, microglial RI was reduced in the striatum of male and female 15-month-old rats compared to both 3- and 9-month-old rats (Fig. 2E; Kruskal-Wallis One-Way: Male: H_(2)_ = 36.25, *p* < .0001, η^2^ = 0.048; 3 vs 15: *p*_dunn_ < .0001; 9 vs 15: *p_dunn_* < .0001; Female: H_(2)_ = 64.53, *p* < .0001, η^2^ = 0.098; 3 vs 15: *p*_dunn_ < .0001; 9 vs 15: *p_dunn_* < .0001). Overall, this analysis revealed significant group-level differences in microglial morphology and highlighted the wide range of morphologies present within each group.

**Figure 2.**
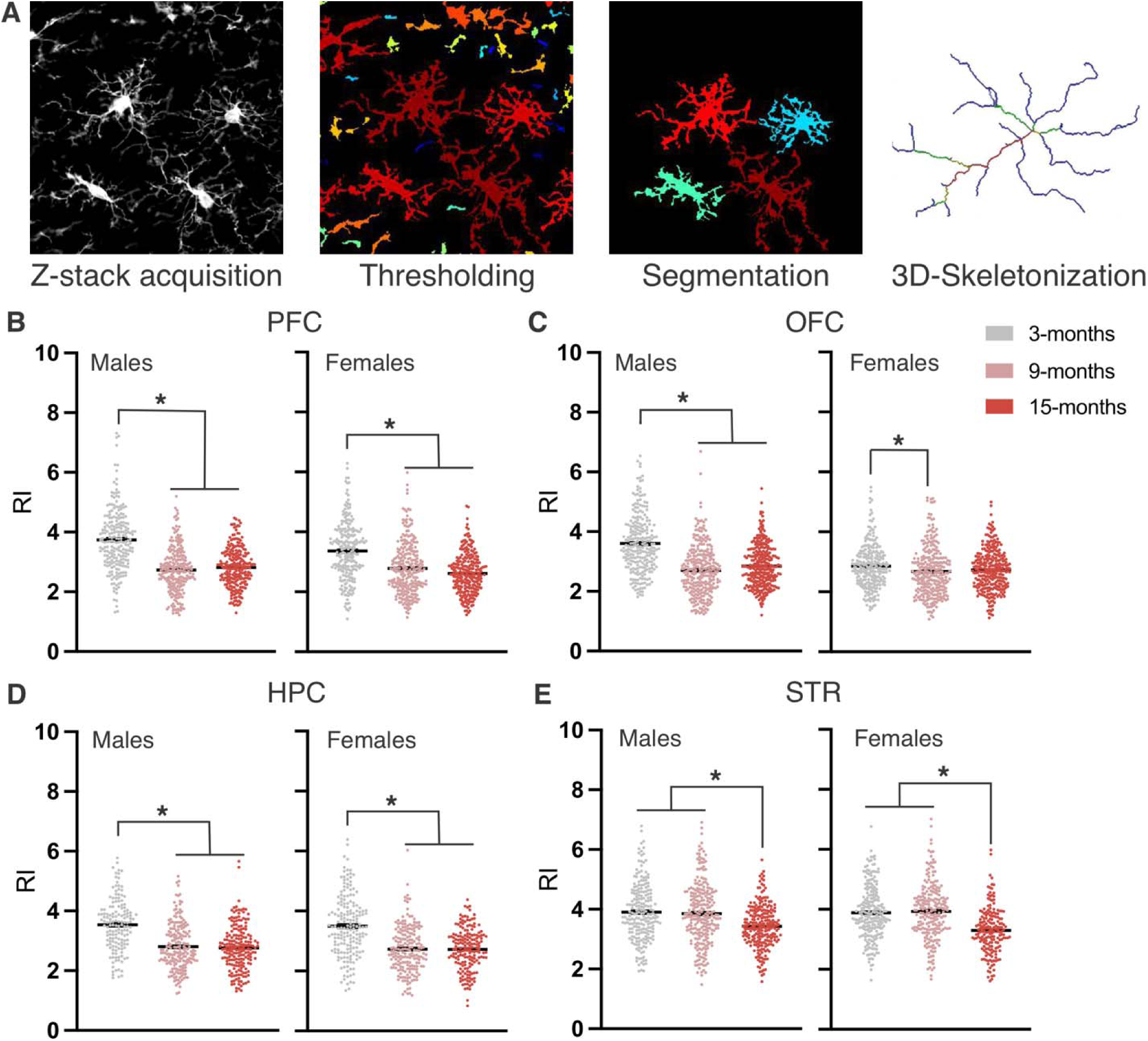
3D morphological assessment shows age-related changes in ramification index. A) 3D Morph analysis workflow. B-E) Ramification index of each cell in 3-, 9-, and 15-month male and female WT rats in the PFC, OFC, HPC, and STR. * Indicates statistical significance (*p* < 0.05) measured using the Kruskal-Wallis non-parametric test. Data represent the group mean ± SEM. Data are shown from 150-304 cells/group, obtained from 5 animals per group.

### 3.3 Hierarchical clustering on principal components revealed age- and region-specific changes in microglial cluster abundance

To capture the heterogeneity in microglial morphology and enable animal-specific microglial analyses, we applied unsupervised HCPC to identify distinct microglial morphotypes. To condense the data into a smaller set of variables, we used principal component analysis (PCA) of eight morphological metrics (cell territory, cell volume, ramification index, branchpoints, endpoints, average branch length, maximum branch length, minimum branch length) across all 5580 analyzed cells. PCA lead to two PCs with an eigenvalue > 1 and accounted for 80.55% of the total variance in the dataset (Table 1, Fig. 3A). The coefficients of each of the morphology metrics for PC1 and PC2 are depicted in Table 2 and represent the contribution of each metric to the PCs. Hierarchical clustering of the 5580 microglia based on the first two PCs resulted in three main clusters (Fig. 3B, C). Cluster 1 (C1) included large microglia with elaborate branching, cluster 2 (C2) represented medium-sized microglia with reduced branching, and cluster 3 (C3) encompassed small microglia with short and limited branching (Fig. 3E, Supplementary Fig. 1). To illustrate the distribution of microglial clusters in each region at each age, we determined the percentage of cells in each cluster (Fig. 3D). Next, to assess both group-level differences and generate individual animal values, Z-scores for relative cluster abundance were calculated for each cluster in each region of interest. In the prefrontal cortex, C3 was increased in 9-month male rats, whereas C2 was decreased in 9- and 15-month female rats (Fig. 4A; One-way ANOVA: Male C3: F_(2, 12)_ = 4.57, *p* = 0.034, η^2^ = 0.43; 3 vs 9: *p*_bonf_ = 0.032; Female C2: F_(2, 12)_ = 11.67, *p* = 0.0015, η^2^ = 0.66; 3 vs 9: *p*_bonf_ = 0.0053; 3 vs 15: *p*_bonf_ *=* 0.0028).

**Table 1.** Principal component eigenvalues. PCs with eigenvalues > 1 (PC1 and PC2, highlighted in blue) were retained for clustering.

| PC | Eigenvalue | Variance (%) |
| --- | --- | --- |
| 1 | 5.25 | 65.57 |
| 2 | 1.20 | 14.98 |
| 3 | 0.71 | 8.92 |
| 4 | 0.43 | 5.36 |
| 5 | 0.27 | 3.34 |
| 6 | 0.075 | 0.94 |
| 7 | 0.049 | 0.61 |
| 8 | 0.023 | 0.28 |

**Figure 3.**
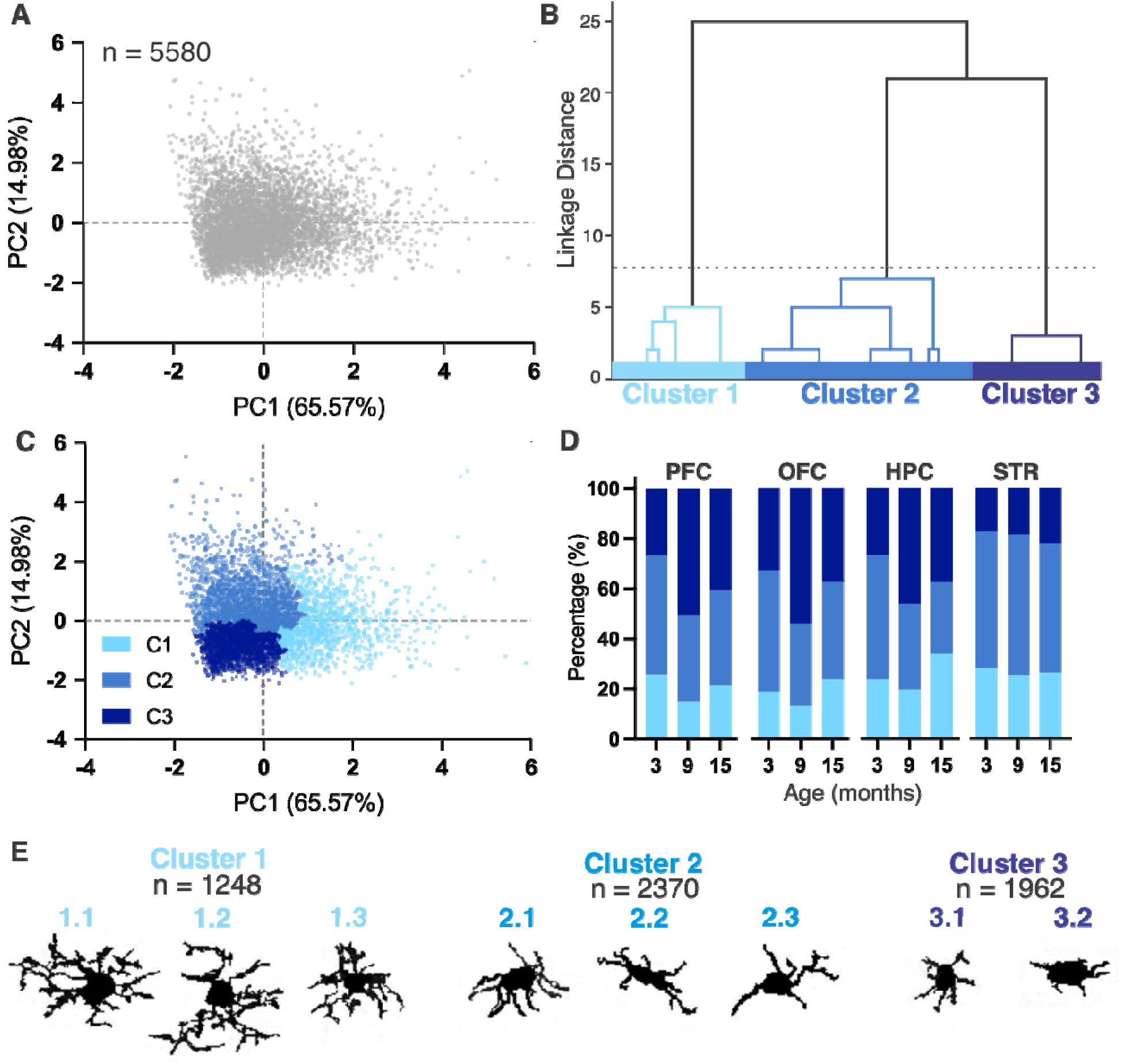
Hierarchical clustering on principal components analysis reveals 3 distinct microglial clusters. A) PCA score plot of all cells on the first two principal components (PC1/PC2) based on 8 morphometric features per cell. B) Hierarchical clustering on PC1/PC2 yielded 3 distinct microglial clusters. C) PCA score plot colour-coded for the three microglial clusters. D) Percentages of each cluster in region of interest in 3-, 9-, and 15-month-old rats. E) Representative cell silhouettes of microglia in each cluster.

**Table 2.**
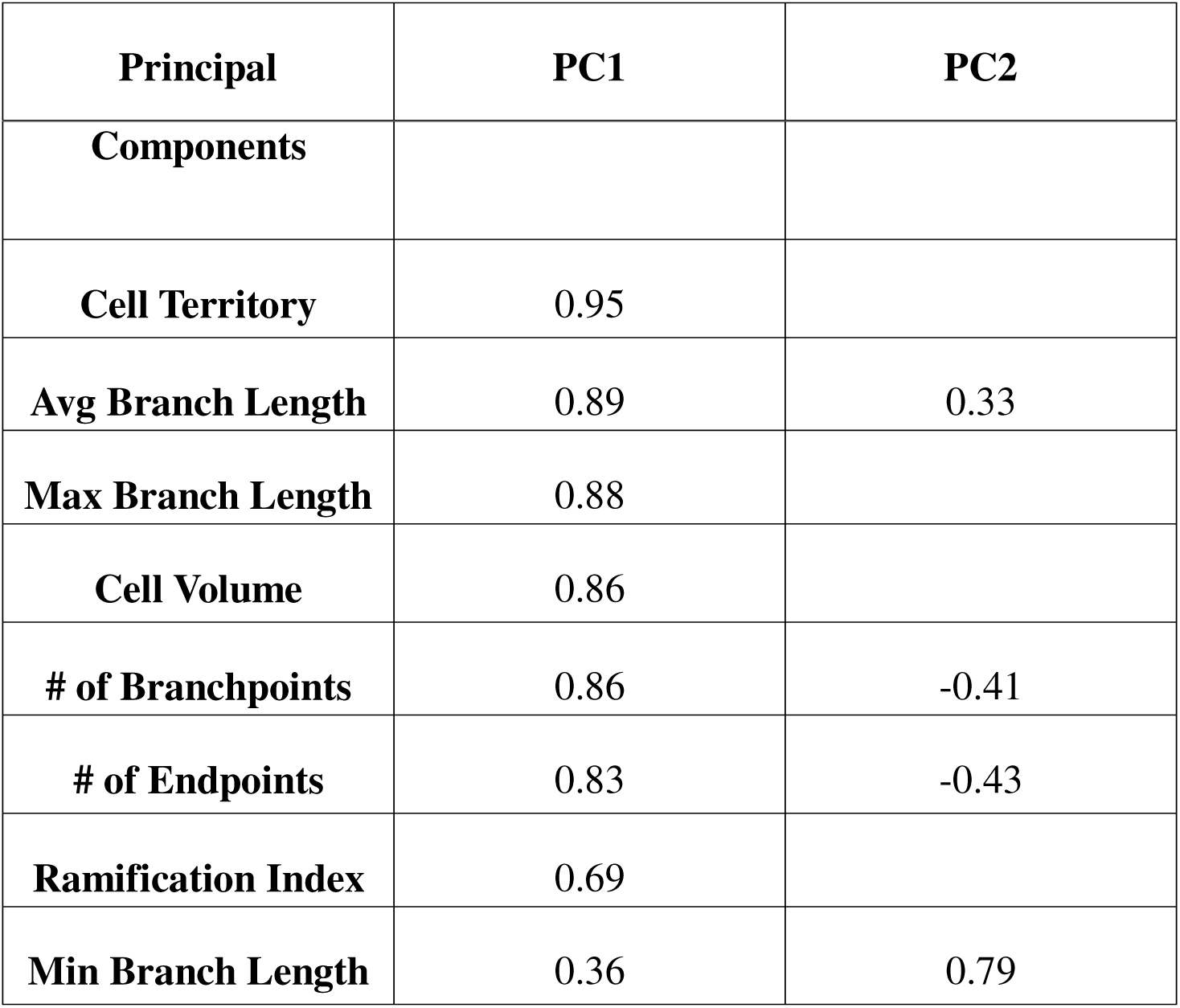
Coefficients of principal component analysis.

**Figure 4.**
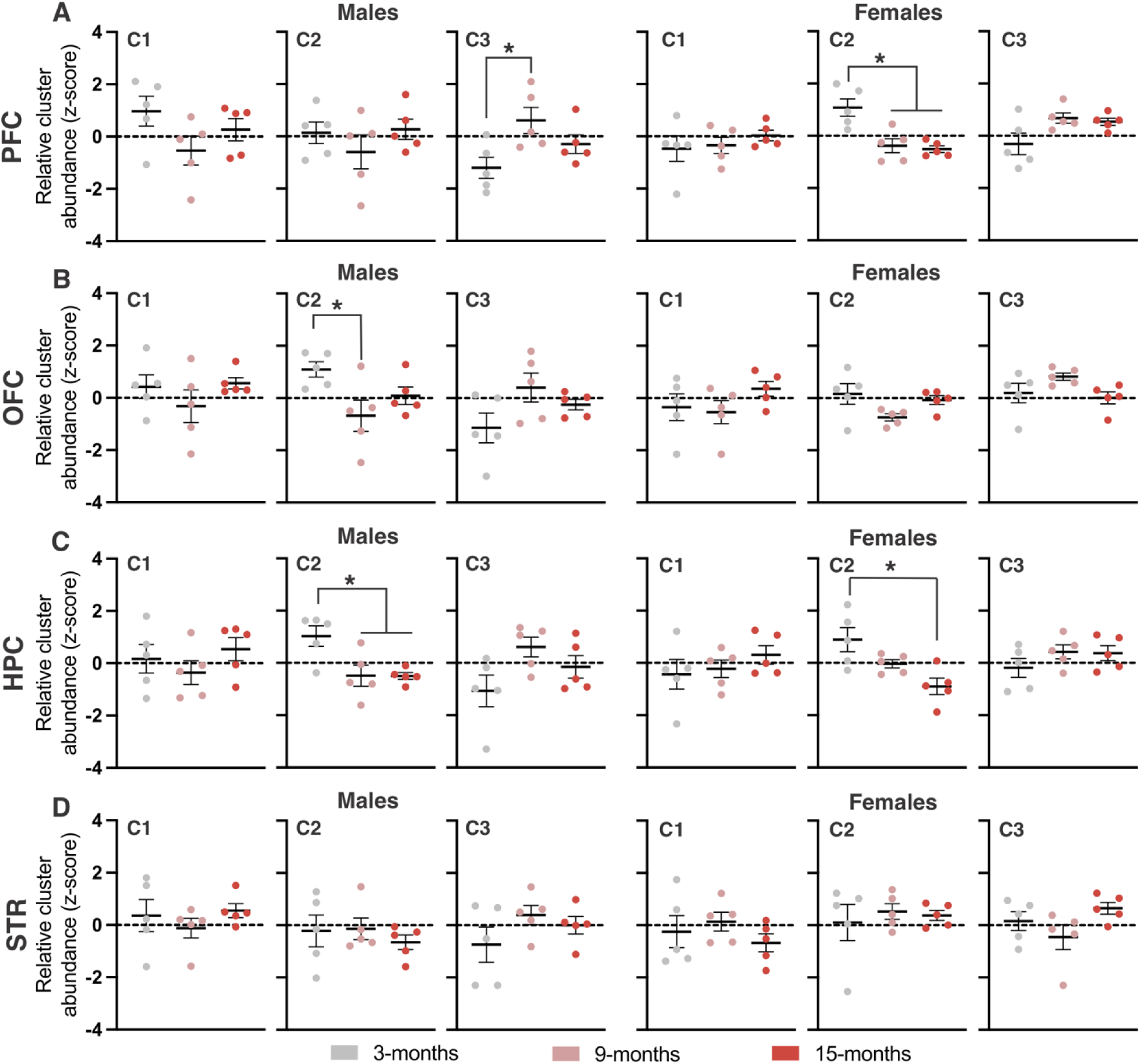
Age-dependent changes in microglial morphotypes. A-D) Z-transformed relative abundance of each cluster in 3-, 9- and 15-month male and female rats. Depicted are A) PFC, B) OFC, C) HPC, and D) STR. * Indicates statistical significance (*p* < 0.05) measured using a one-way ANOVA. Data represent the group mean ± SEM. *n* = 5 males and 5 females per age.

In the orbitofrontal cortex, C2 abundance was reduced in 9-month males, with no significant differences observed in females at any age (Fig. 4B; One-way ANOVA: Male C2: F_(2, 12)_ = 4.21, *p* = 0.041, η^2^ = 0.41; 3 vs 9: *p*_bonf_ = 0.041). In the hippocampus, C2 was significantly reduced in 9- and 15-month males, as well as in 15-month females (Fig. 4C; One-way ANOVA: Male C2: F_(2, 12)_ = 7.082, *p* = 0.0093, η^2^ = 0.54; 3 vs 9: *p*_bonf_ = 0.021; 3 vs 15: *p*_bonf_ *=* 0.020; Female C2: F_(2, 12)_ = 7.138, *p* = 0.0091, η^2^ = 0.54; 3 vs 15: *p*_bonf_ = 0.0079). Finally, no significant age-related differences in cluster abundance were observed in the striatum (Fig. 4D). Altogether, we used HCPC to reveal three distinct clusters of microglia and changes in their relative abundance across age.

### 3.4 Spatial reference and working memory was impaired in 9- and 15-month-old rats

To determine the effects of age on spatial working and reference memory, we employed a 4/8 radial arm water maze (RAWM) task. Errors were summed in three-day blocks for both reference and working memory to evaluate performance over time (Fig. 5B, C). Rats of all ages exhibited spatial learning, as indicated by a progressive decrease in errors across the 15 testing days for both reference (Fig. 5B) and working memory (Fig 5C). Age-related impairments in spatial reference memory (entries down arms that never contained a platform) varied across testing days. In males, 9-month-old rats committed more errors than 3-month-old rats on day 6, while both 9-and 15-month-old rats made more errors than 3-month-old rats on days 9 and 12 (Fig. 5B; Two-way mixed ANOVA: Male: Day x Age interaction; F_(8, 108)_ = 2.59, *p* = 0.013, η ^2^ = 0.16; Day 6: 3 vs 9: *p*_bonf_ = 0.028; Day 9: 3 vs 9: *p*_bonf_ = 0.017, 3 vs 15: *p*_bonf_ = 0.002; Day 12: 3 vs 9: *p*_bonf_ < .001, 3 vs 15: *p*_bonf_ = 0.015). In female rats, 15-month rats performed worse than 3-month rats on day 6, and 9-month-old rats committed more errors than 3-month-old rats on day 15 (Fig. 5; Two-way mixed ANOVA: Female: Day x Age interaction; F_(8, 108)_ = 2.77, *p* = 0.008, η ^2^ = 0.17; Day 6: 3 vs 15: *p*_bonf_ = 0.005; Day 15: 3 vs 9: *p*_bonf_ = 0.11). To further evaluate spatial reference memory, we analyzed errors across individual trials and total errors accumulated over the entire testing period (Fig. 5D, E). Nine-and 15-month-old males committed more reference memory errors overall compared to 3-month-old males (Fig. 5E; One-way ANOVA: Male: F_(2, 29)_ = 15.18, *p* < .001, η^2^ = 0.53; 3 vs 9: *p*_bonf_ < .001; 3 vs 15: *p*_bonf_ = 0.001) with specific increases on trials 2, 3, and 4 (Fig. 5D; Two-way mixed ANOVA: Male: Trial x Age interaction; F_(4.3, 58.3)_ = 3.019, *p* = 0.022, η ^2^ = 0.18; Trial 2: 3 vs 9: *p*_bonf_ = 0.005, 3 vs 15: *p*_bonf_ = 0.002; Trial 3: 3 vs 9: *p*_bonf_ < .001, 3 vs 15: *p*_bonf_ = 0.003; Trial 4: 3 vs 9: *p*_bonf_ = 0.012). In contrast, no age-related differences in total reference memory errors (Fig. 5E) or trial-specific performance was observed in females (Fig. 5D). Spatial working memory (entries down arms that previously contained platforms) deficits in males also varied by testing day. Nine-month males made more errors than 15-month males on day 3, performed worse than 3-month males on days 9 and 15, and both 9- and 15-month males committed more errors than 3-month males on day 12 (Fig. 5C; Two-way mixed ANOVA: Male: Day x Age interaction: F_(6.1, 82.9)_ = 2.44, *p* = 0.031, η ^2^ = 0.15; Day 3: 9 vs 15: *p*_bonf_ = 0.011; Day 9: 3 vs 9: *p*_bonf_ = 0.002; Day 12: 3 vs 9: *p*_bonf_ = 0.005, 3 vs 15: *p*_bonf_ = 0.04; Day 15: 3 vs 9: *p*_bonf_ = 0.006). In females, both 9- and 15-month rats made more working memory errors than 3-month rats across all testing days (Fig. 5C; Two-way mixed ANOVA: Female: main effect of age; F_(2, 27)_ = 6.003, *p* = 0.007, η ^2^ = 0.31; 3 vs 9: *p*_bonf_ = 0.013; 3 vs 15: *p*_bonf_ = 0.023). Further analysis of total working memory errors showed that 9-month males committed more errors than both 3- and 15-month males overall (Fig. 5G; One-way ANOVA: Male: F_(2, 29)_ = 10.11, *p* < .001, η^2^ = 0.43; 3 vs 9: *p*_bonf_ < .001; 9 vs 15: *p*_bonf_ = 0.037) and across trials (Fig. 5F; Two-way mixed ANOVA: Male: main effect of age; F_(2, 27)_ = 10.11, *p* < .001, η ^2^ = 0.43; 3 vs 9: *p*_bonf_ < .001; 9 vs 15: *p*_bonf_ = 0.037). Similarly, both 9- and 15-month females committed more total working memory errors than 3-month females (Fig. 5G; One-way ANOVA: Female: F_(2, 29)_ = 6.003, *p* = 0.007, η^2^ = 0.31; 3 vs 9: *p*_bonf_ = 0.013; 3 vs 15: *p*_bonf_ = 0.023) independent of trial (Fig. 5F; Two-way mixed ANOVA: Female: main effect of age; F_(2, 27)_ = 6.37, *p* = 0.005, η ^2^ = 0.32; 3 vs 9: *p*_bonf_ = 0.011; 3 vs 15: *p*_bonf_ = 0.018). Together, these findings highlight nuanced, age- and sex-specific patterns of spatial reference and working memory performance in rats.

**Figure 5.**
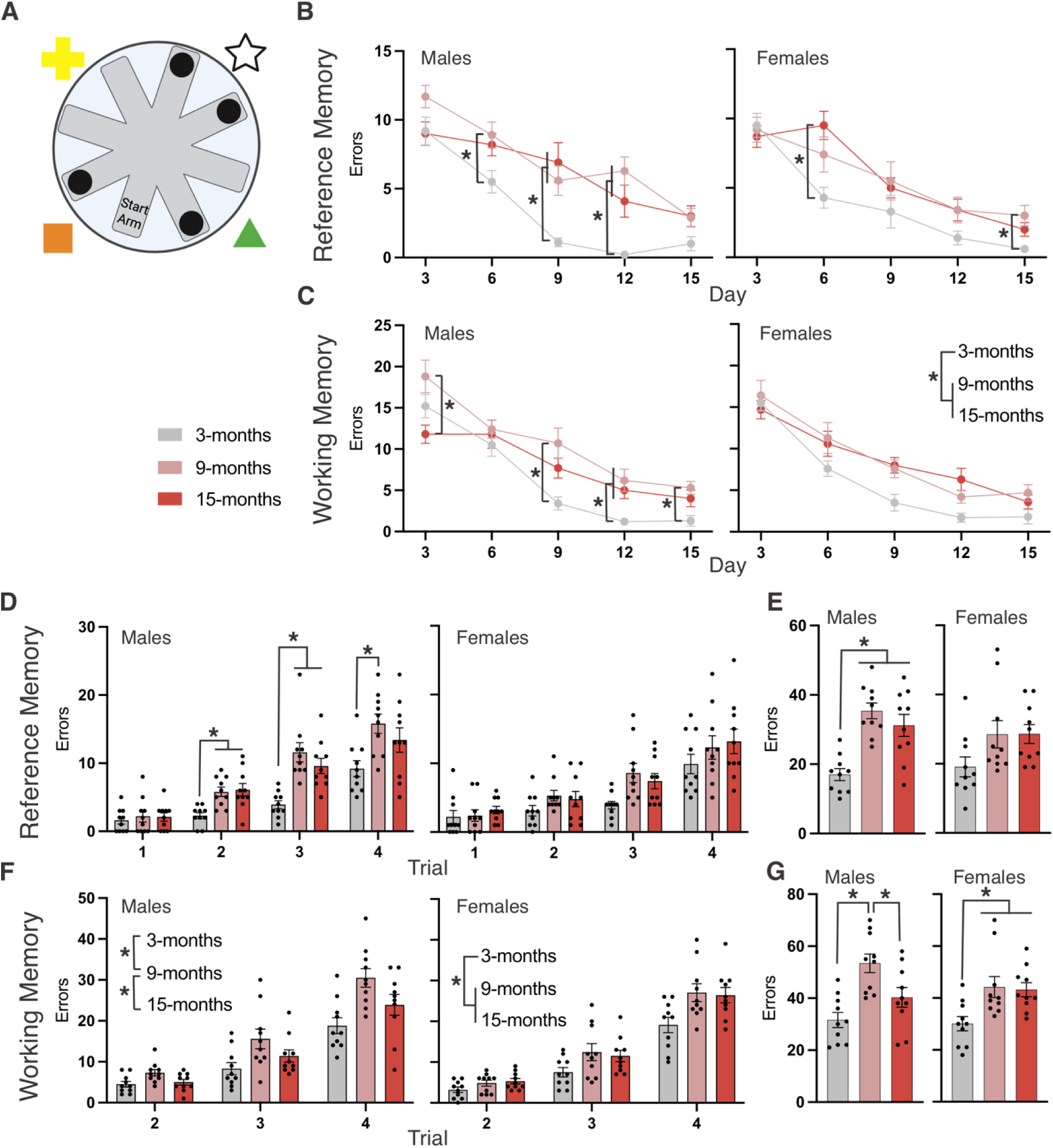
Impaired spatial reference and working memory in aged rats. A) Schematic of RAWM maze and platform setup. B) Reference memory errors committed across the 15 testing days displayed as sums per 3-day increment in 3-, 9-, and 15-month WT rats. C) Working memory errors committed across the 15 testing days displayed as sums per 3-day increment in 3-, 9-, and 15-month WT rats. D) Total reference memory errors committed in each trial. E) Total reference memory errors committed across all 15 testing days. F) Total working memory errors committed in each trial. G) Total working memory errors committed across all 15 testing days. * Indicates statistical significance (*p* < 0.05). Data represent the group mean ± SEM. *n* = 10 males and 10 females per group.

### 3.5 Set-shifting performance was mainly unaffected by age, except 9-month female rats performed worse in response discrimination

In addition to spatial memory, we evaluated striatal-dependent learning and cognitive flexibility though a water-based T-maze set-shifting task. First, rats underwent visual cue discrimination (VCD); a striatal-driven task in which they learned to navigate to the arm with an illuminated light to locate the hidden platform (Fig. 6A). Performance was quantified by the number of trials required to reach a rolling average of 10 out of 12 correct responses, as well as by the total number of errors. No significant age-related deficits in VCD were observed in either male or female rats (Fig. 6B, C). Upon reaching criterion in VCD, the task shifted to response discrimination (RD). In this stage, the light continued to alternate pseudo-randomly, but the hidden platform stayed fixed on one side of the maze (Fig. 6D). While male rats showed no age-related RD deficits, 9-month-old females exhibited impairments, requiring more trials (Fig. 6E; One-way ANOVA: Female: F_(2, 29)_ = 5.47, *p* = 0.01, η^2^ = 0.29; 3 vs 9: *p*_bonf_ = 0.17; 9 vs 15: *p*_bonf_ = 0.36) and committing more errors than both 3- and 15-month females (Fig. 6F; One-way ANOVA: Female: F_(2, 29)_ = 14.38, *p* < .001, η^2^ = 0.52; 3 vs 9: *p*_bonf_ < .001; 9 vs 15: *p*_bonf_ < .001).

**Figure 6.**
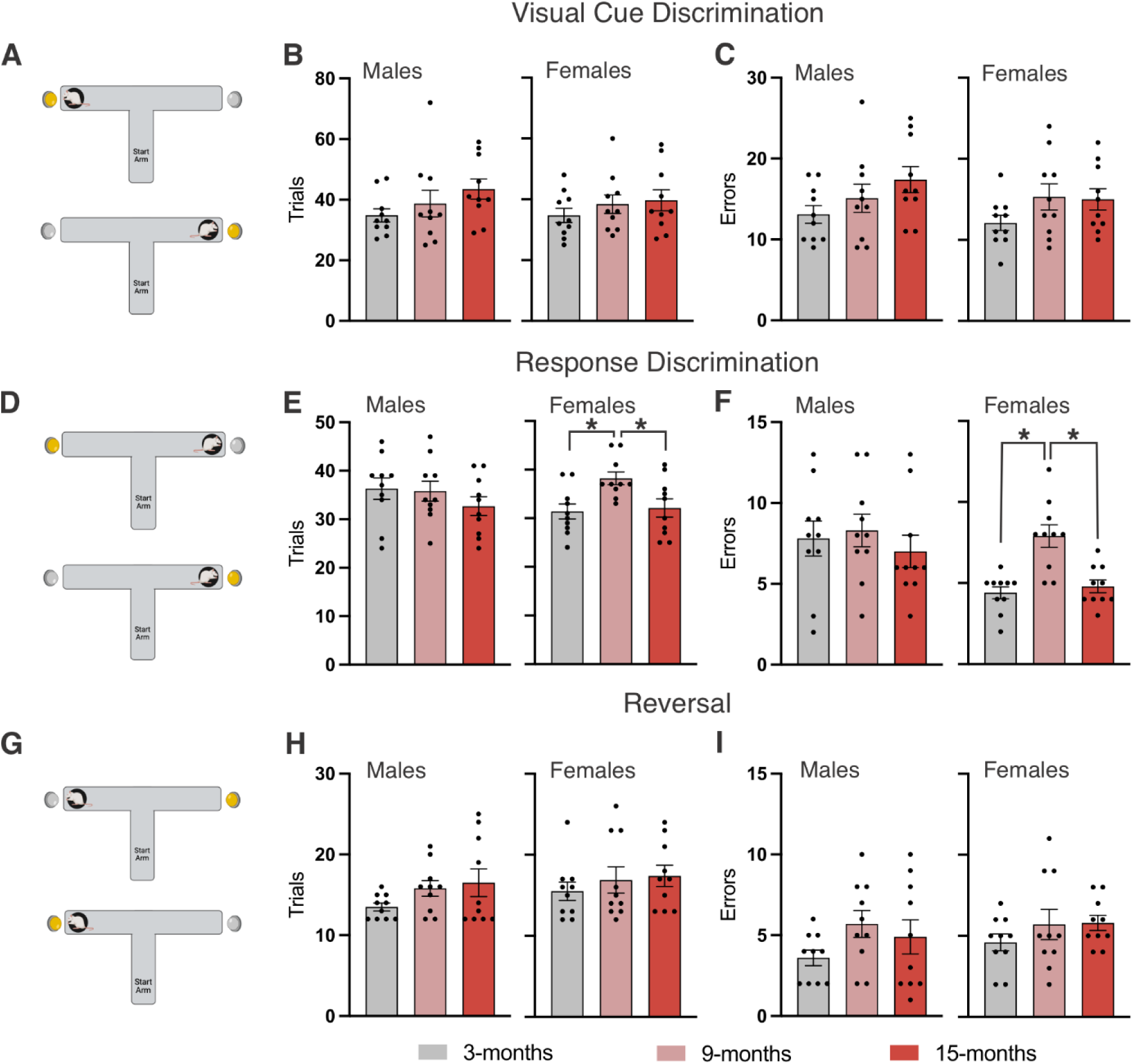
Female-specific age-related impairment in response discrimination, with visual cue discrimination and reversal learning unaltered. A) The rats learned to associate the illuminated light with the escape platform in the visual cue discrimination (VCD) phase of the task. B) Trials taken to reach a 10/12 rolling average in VCD. C) Errors committed prior to completing VCD phase of the task. D) Cognitive flexibility was assessed by shifting the rule to “stay to one side” and ignore the light in the response discrimination (RD) portion of the task. E) Trials taken to reach a 10/12 rolling average in incongruent trials in RD. F) Errors committed prior to completing the RD phase of the task. G) Cognitive flexibility was assessed again by switching the platform to the other side of the T-maze for a reversal. H) Trials taken to reach a 10/12 rolling average in reversal. I) Errors committed prior to completing the reversal portion of the task. * Indicates statistical significance (*p* < 0.05) measured using a one-way ANOVA. Data represent the group mean ± SEM. *n* = 10 males and 10 females per age.

Once criterion was achieved in RD, rats progressed to reversal learning (RL), in which the platform location was switched to the opposite arm, but the task remained otherwise identical (Fig. 5G). No significant differences were observed in the RL phase of testing. Although group-level differences were limited across phases, these results revealed variability in individual performance that may relate to region-specific microglial morphotypes.

### 3.6 Cognitive performance correlated with microglia cluster abundance in relevant brain regions

To comprehensively explore potential relationships between microglial morphotypes and cognition across all rats, we generated a correlation matrix encompassing behavioural measures and region-specific microglial cluster abundances (Supplementary Fig. 2). Several notable correlations emerged between microglial morphotypes and performance, particularly in brain regions implicated in the corresponding cognitive domains. In the prefrontal cortex, C2 microglial abundance was negatively associated with response discrimination errors, such that lower C2 levels were linked to poorer performance (Fig. 7A; *p* = 0.036, R^2^ = 0.15). In the orbitofrontal cortex, C1 abundance was negatively associated with reversal errors, whereas C3 abundance showed a positive correlation with reversal errors (Fig. 7B; C1: *p* = 0.0045, R^2^ = 0.25; C3: *p* = 0.0017, R^2^ = 0.30). Similarly, in the hippocampus, C2 abundance was negatively associated with total reference memory errors (Fig. 7C; *p* = 0.031, R^2^ = 0.15). These correlations highlight the potential role of region-specific microglial morphotypes as contributors to, or indicators of, cognitive performance in aging.

**Figure 7.**
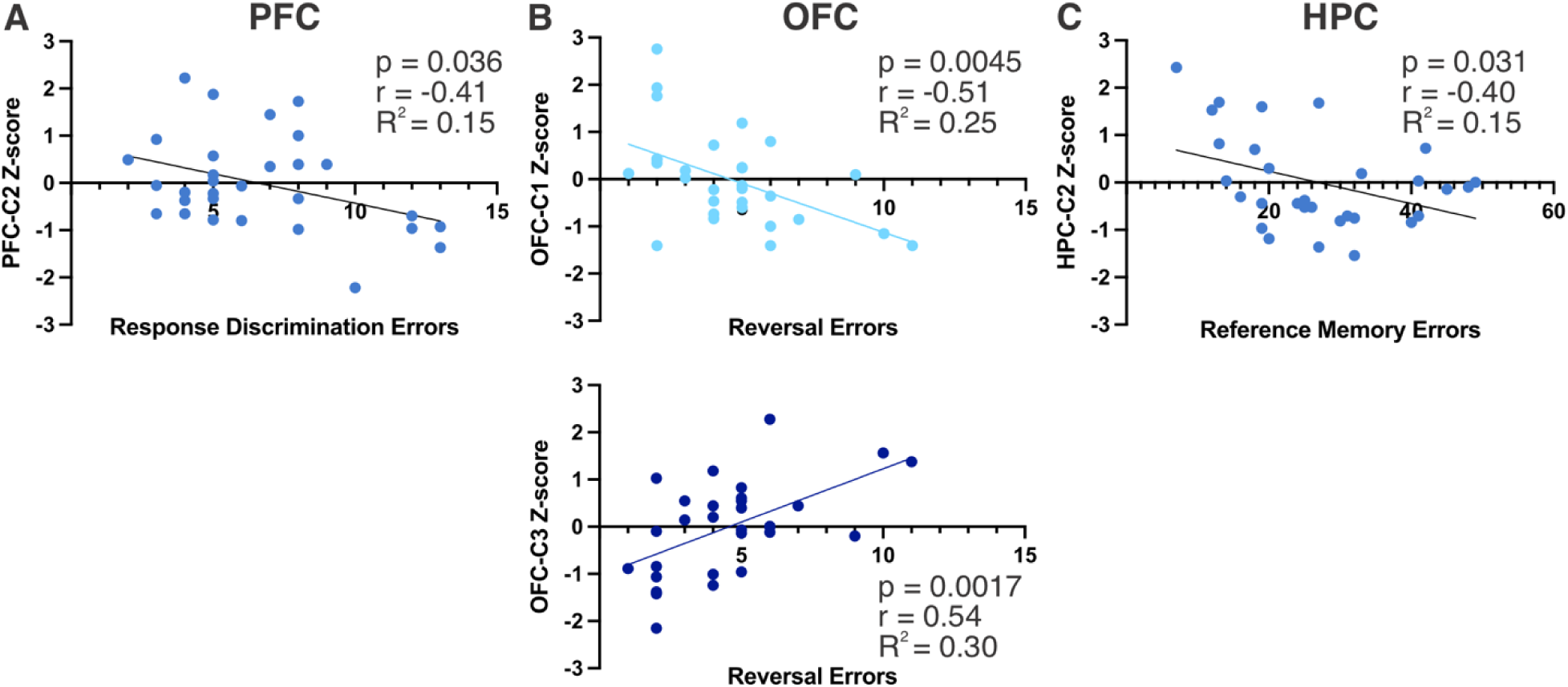
Region-specific morphotype abundance correlates with cognitive outcomes. A) Simple linear regression between C2 PFC microglia and response discrimination errors. B) Simple linear regressions between C1 OFC microglia and reversal errors as well as C3 OFC microglia and reversal errors. C) Simple linear regression between C2 HPC microglia and reference memory errors. *n* = 10 males and 10 females per group.

## 4. Discussion

This study represents the first comprehensive investigation linking region-specific microglial morphological phenotypes to cognitive performance in normal aging. Using an innovative unsupervised clustering approach, we demonstrate for the first time that distinct microglial morphotypes emerge with aging and that their regional abundance directly correlates with cognitive function in corresponding brain regions. Our findings reveal a previously unrecognized relationship between microglial structural diversity and cognitive aging, showing that age-related cognitive decline is not uniform across domains. Spatial reference memory, spatial working memory, and cognitive flexibility were impaired to varying degrees, while striatal-based learning remained intact. Most importantly, we discovered that specific microglial morphotypes in functionally relevant brain regions serve as cellular correlates of cognitive performance: prefrontal cortex microglia correlated with executive function, orbitofrontal cortex microglia associated with behavioral flexibility, and hippocampal microglia correlated with spatial reference memory. This work establishes, for the first time, that microglial morphological heterogeneity is not merely a consequence of aging but represents a meaningful cellular signature of cognitive status in the aging brain. These findings advance our understanding of how microglia contribute to cognitive aging and provide a novel framework for investigating microglia-cognition relationships in health and disease.

### 4.1 Age-related alterations in microglial cell density

Although microglial morphology was the primary focus of this study, we also examined whether microglial cell density varied by age in our regions of interest. Previous studies report mixed findings on age-related changes in microglial density, with effects differing by brain region and model. Increases in microglial density have been observed in the aged mouse hippocampus, thalamus, and sensory cortices (visual and auditory) ^40–42^. However, a study in aged rats reported no change in hippocampal microglial density with age^43^. Similarly, clinical studies have reported no change in microglial density in cortical grey matter with aging, despite changes in morphology^27,42^. Consistent with these findings, we observed no age-related differences in microglial cell density in the prefrontal cortex, orbitofrontal cortex, or hippocampus (Fig. 1C, F, I). However, a reduction in microglial density was observed in the striatum of 15-month-old females (Fig. 1L), in line with previous work describing reduced microglia numbers in the aged mouse striatum^44^. Altogether, our results reinforce the notion that age-dependent changes in microglial cell numbers vary by anatomical region and that morphological changes can occur independently of cell number.

### 4.2 Region-specific changes in microglial morphotype abundance across age

To capture how microglial aging may differ throughout the brain, we quantified individual morphometric features in four cognitive brain regions. Previous work has shown that aging is associated with reduced microglial branching in both rodent and human studies, particularly in the hippocampus and cortical regions^26,27,45,46^. Consistent with previous reports, and expanding on the anatomical scope, we observed age-related reductions in ramification index by 9-months of age in the prefrontal cortex, orbitofrontal cortex, and hippocampus (Fig. 2B, C, D).

Ramification index was also reduced in the striatum, but only in 15-month-old rats (Fig. 2E), highlighting that age-related morphological changes are not uniform throughout the brain. While this approach captured meaningful age-related differences, it didn’t identify distinct structural phenotypes or produce values suitable for correlational analyses.

Prior studies examining microglial morphology across age have often relied on qualitative descriptions or predefined, subjective categorizations of microglial types. In contrast to this, and to better capture microglial heterogeneity, we applied hierarchical clustering on principal components to our morphological metrics^47^. This unsupervised approach allowed us to classify microglia into distinct morphotypes based on shared morphological features. The resulting three clusters differed in cell size, branching complexity, and branch length (Fig. S1). We observed region-specific shifts in relative cluster abundance across age. For example, significant age-related decreases in C2 (medium-sized microglia with reduced branching) were observed in the female prefrontal cortex, the male orbitofrontal cortex, and both the male and female hippocampus (Fig. 4A-C). In addition, C3 (small microglia with short and limited branching) was increased in the prefrontal cortex of 9-month-old males (Fig. 4A). Interestingly, despite an overall increase in ramification index in 15-month-old rats, no differences in cluster abundance were observed in the striatum (Fig. 4D). These findings suggest that aging drives differential microglial responses depending on anatomical region and sex.

Given our sample size, we analyzed age effects separately for each sex to preserve statistical power. This approach revealed that aging affected microglial cluster abundance differently in males versus females across brain regions, suggesting sex-specific patterns of microglial aging. These observations are consistent with growing evidence from both human and rodent studies describing significant differences in the molecular profiles of male and female microglia^48–52^. For example, female microglia have been described as having heightened surveillance and reactivity in rodents and similarly have elevated inflammatory gene expression profiles in humans^50,52^. Although the functional consequences of this remain unclear, it highlights the importance of inclusion of sex as a factor in future studies of microglia and aging.

### 4.3 Correlating microglial morphology with cognitive function

Having established region- and age-specific changes in microglial morphology, our next objective was to examine whether these changes corresponded with cognitive performance. While age-related impairments in reference memory have been well described, executive function (i.e., working memory, cognitive flexibility) remains relatively understudied in rodent models. Given that aging can impact a range of domains^3–6^, including executive function, we employed two behavioural tasks to capture changes in spatial working memory, spatial reference memory, striatal-based learning, and cognitive flexibility. In line with previous reports of reference memory decline in aged rodents^4,53,54^, we observed impairments in the spatial reference memory component of the RAWM task in 9- and 15-month-old male rats, and to a lesser extent in females (Fig. 5B, D, E). Similarly, our results support previous evidence of working memory decline with age in rodent models^54–56^, as we found female 9- and 15-month rats were impaired in the spatial working memory component of the RAWM task (Fig. 5C, F, G). Interestingly, male 9-month-old rats committed more working memory errors than younger or older males, which may reflect differences in strategy use or a shift in regional circuitry reliance at midlife (Fig. 5F, G). These results confirm that age-related cognitive impairments may vary by sex and cognitive domain and can manifest mid-adulthood.

We next evaluated learning and cognitive flexibility using a set-shifting task involving visual and directional discrimination. Striatal-based learning, assessed via visual cue discrimination, did not differ significantly across ages (Fig. 6A-C), consistent with previous work from our lab that demonstrated preserved visual cue discrimination in a lever-pressing task at 8- and 13-months of age^4^. However, we observed increased trials and errors in the response discrimination phase of testing in 9-month-old females (Fig. 6D-F). This finding was somewhat unexpected given the lack of impairment at 15 months and could reflect early disruptions in flexibility or inhibitory control that may not progress linearly with age. Notably, we observed substantial variability in performance across all phases of the task suggesting individual differences in aging trajectories. The variability in our rats is consistent with prior reports of significant cognitive heterogeneity in mouse and rat models of aging^55,57,58^.

Previous studies have observed a relationship between reactive microglial markers and cognition in models of aging and AD^29–33^. Based on this, we hypothesized that microglia morphology in cognitive brain regions might also associate with age-related cognitive decline. Given the substantial variability in performance in each group, and lack of consistent group-level differences, we collapsed animals across age and sex for correlational analyses. We found that microglial cluster abundance was selectively associated with cognitive performance in regionally relevant cognitive domains. Specifically, in the prefrontal cortex and hippocampus, regions critically important in executive function and spatial memory, higher abundance of C2 negatively correlated with response discrimination and reference memory errors respectively (Fig. 7A, C). In contrast to this, C1 and C3 in the orbitofrontal cortex, a region implicated in flexibility and reversal, were significantly correlated with errors committed in reversal learning (Fig. 7B). Although we cannot determine whether these microglial features are causative or reflective of age-related cognitive impairment, these results suggest that different microglial morphotypes may reflect regionally distinct aging processes with functional consequences.

### 4.4 Experimental limitations and future directions

An important consideration in interpreting these results is that microglia are highly active and can rapidly shift their phenotypes in response to environmental cues. Our analysis captures these cells at a single point in time, providing a snapshot of cluster distributions rather than a full view of the microglial dynamics. Future work incorporating live-imaging approaches may add to our understanding of microglial morphology across aging. In addition, although we identified clusters based on shared morphometric features, we did not attempt to classify or name these clusters. Based mostly on human literature, dystrophic microglia are an important aging phenotype that our analysis did not account for^26,27^. The future inclusion of specific features like process bulbs or fragmentation may help clarify the presence of dystrophic microglia in this model, which may be associated with impaired surveillance and synaptic regulation.

While our findings offer important insight into microglial heterogeneity, they also raise questions about the functional relevance of the identified morphotypes. Future studies should aim to integrate morphometric profiling with transcriptomic and proteomic approaches to better understand the cellular mechanisms underlying cognitive decline and how they relate to morphology. In addition, assessing microglial-synapse interactions could reveal whether certain morphotypes are involved in synapse maintenance or loss, which is among the strongest predictors of cognitive decline in AD^59^. It is also important to acknowledge potential species differences in microglia, as prior studies have shown major differences in transcriptomic profiles of human versus rodent microglia^60^, but further work is needed to consider species-related differences in microglia across age.

In conclusion, we used HCPC to identify unique microglial morphotypes in cognitive brain regions and characterized their changes across age. To our knowledge, this is the first study to apply unsupervised clustering to microglial morphometrics in an aging model, and in the future could serve as a complementary approach to microglial phenotyping being conducted in models of neurodegeneration and injury. We also used a multidomain approach to behavioural testing that allowed us to associate cognitive performance with microglial changes in relevant brain regions. While further work is needed to elucidate the molecular and functional profiles of microglia in relation to their morphology, we provide important evidence for microglial structural changes being a meaningful correlate of cognitive status in normal aging. The combination of microglial morphotyping with molecular phenotyping represents an exciting next direction for furthering our understanding of microglia during aging.

## Ethics approval

Animal ethics and procedures used in this study were approved by the Animal Care Committee at Western University (protocol 2022-137). All rats included in this study were housed in facilities maintained by Western University Animal Care and Veterinary Services.

## Acknowledgements

We would like to thank Dr. Lynn Wang and UWO’s Animal Care and Veterinary Services staff for their technical assistance.

## Data availability

Data will be made available on request.

