## Supplemental Figures for "Microglial morphology reflects cognitive status in the aging rat brain"

Figures: 7

Correspondence:

Shawn Whitehead, PhD

Department of Anatomy and Cell Biology

458 Medical Sciences Building

Western University

London, Ontario

N6A 3K7

519 661-2111 x80440

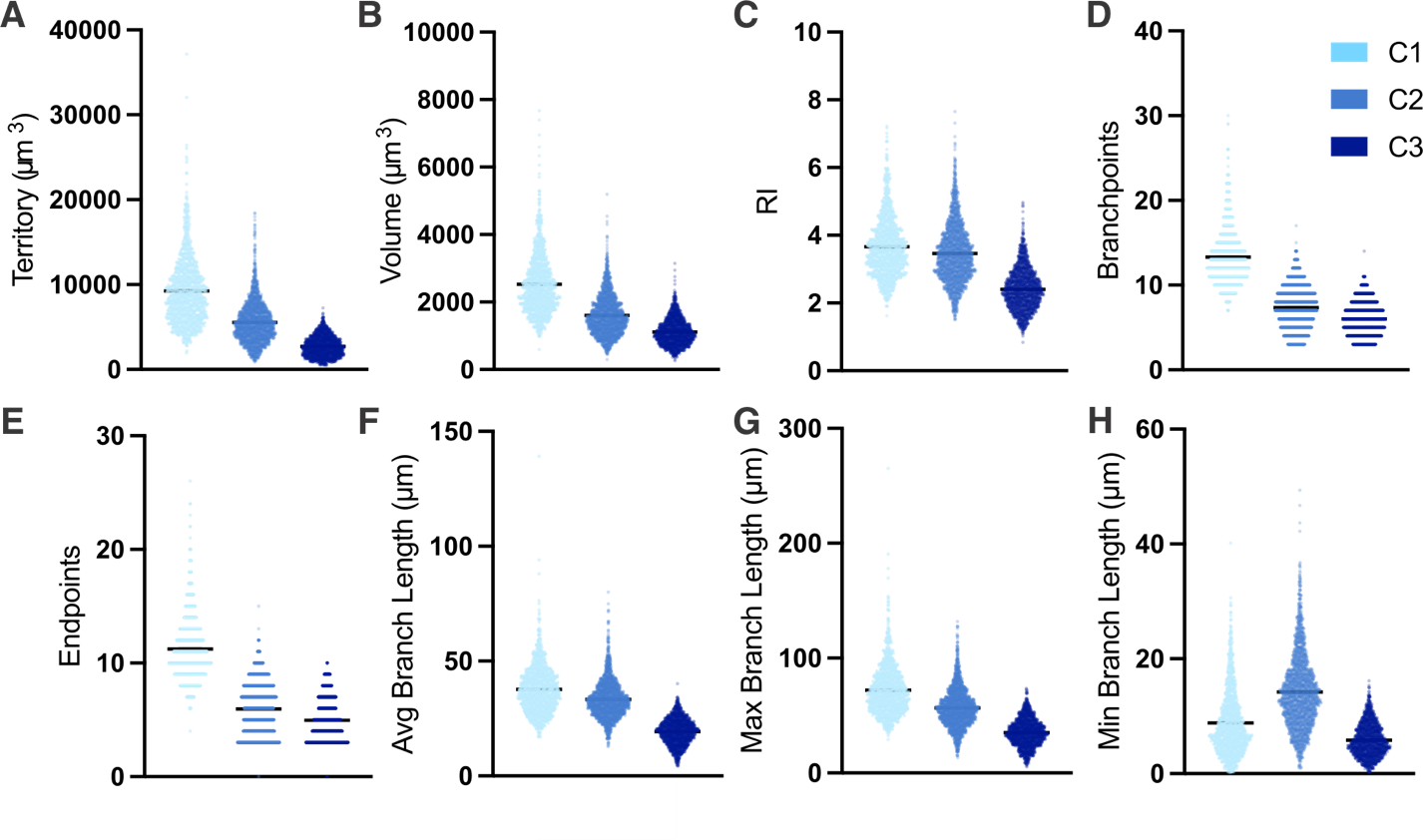
**Supplemental Figure 1. Morphometric features for microglial clusters.** Depicted are A) cell territory, B) cell volume, C) ramification index, D) number of branchpoints, E) number of endpoints, F) average branch length, G) maximum branch length, H) minimum branch length.

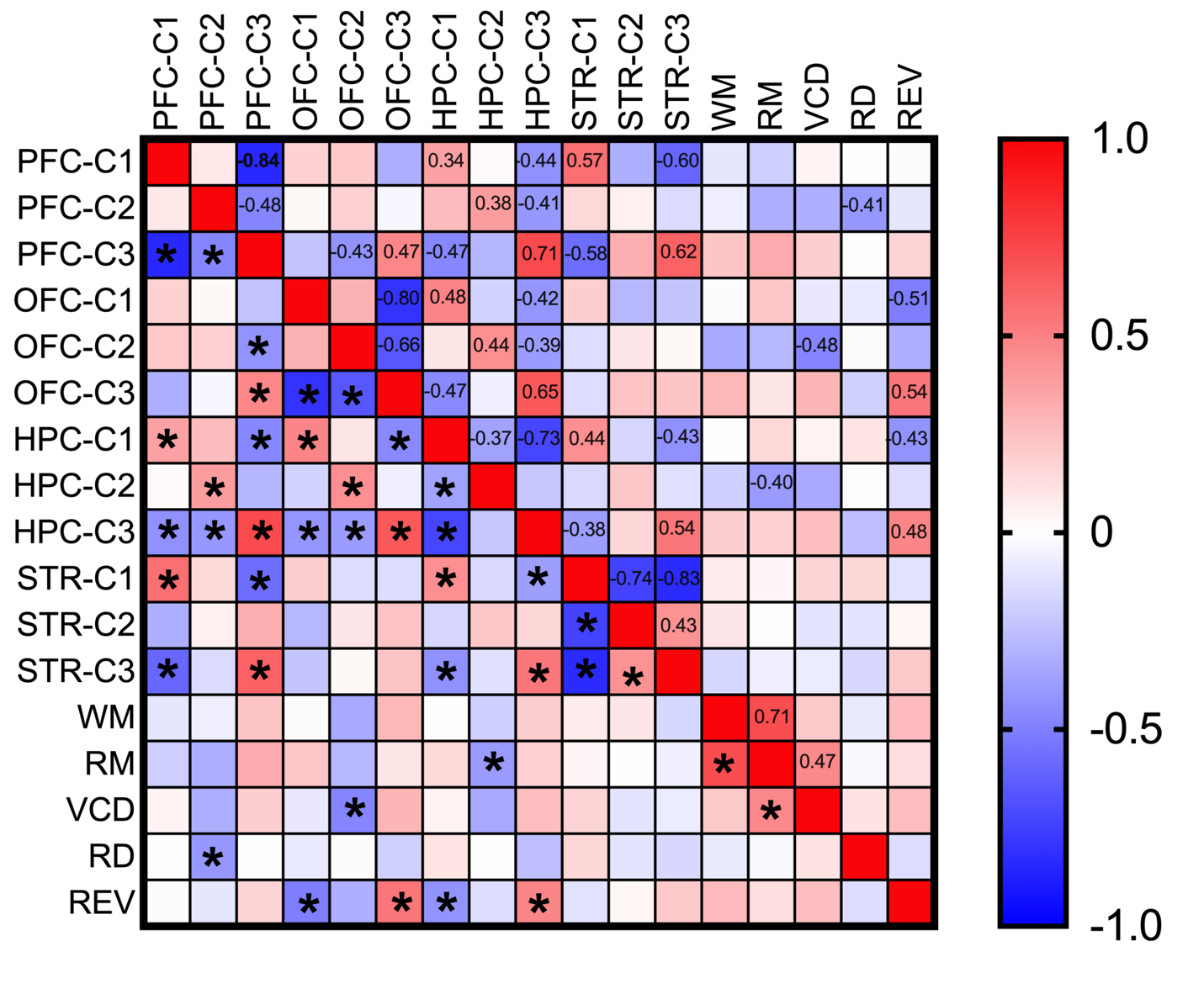
**Supplemental Figure 2.** Pearson correlation matrix showing relationships between performance on cognitive tasks and abundance of microglial morphology clusters in different brain regions. Positive correlations are presented with red shading, and negative correlations are presented with blue shading. Colour intensity corresponds to the r-value and r-value is shown in the upper half of the matrix for significant correlations. * Indicates statistical significance (*p* < 0.05).
